# Recent history attracts and repels perceptual decisions depending on surprise

**DOI:** 10.64898/2026.06.25.734467

**Authors:** Aaron Kaltenmaier, Clare Press

## Abstract

Past sensory experience shapes our perceptual decision-making in the now. Popular models frame perceptual decisions as *either* attracted towards or repelled away from recent sensory information, but it is unclear when and why these distinct effects emerge. We here ask whether effects turn from attractive to repulsive depending on the level of surprise elicited by the precision-weighted discrepancy between past and present sensory states. This model is based upon the idea that attraction is adaptive for optimising efficiency and accuracy when discrepancies are small, because they likely reflect sensory noise rather than real change in the environment. In contrast, repulsion may reflect the upweighting of counterfactual evidence when discrepancies are large because they more likely signal the need for model updating. We test this model on a large amount of recently-collated trial-by-trial serial dependence data and consistently find support for it across the dataset, participant, and trial-by-trial level. Specifically, serial dependence effects are attractive at low discrepancies between past and current sensory states but turn repulsive when discrepancies are larger. Higher sensory precision is found to accelerate this flip by reducing the modal discrepancy threshold required to trigger repulsion effects. We discuss how these findings necessitate extending existing theories of serial dependence, and how they may resolve conflicts in the broader predictive processing, learning and perception literatures.

## Introduction

Perception is increasingly understood as a process of inference, where incoming signals are combined with information from our past (Friston, 2005; Keller & Mrsic-Flogel, 2018). Perception therefore no longer faithfully resembles the incoming evidence as it reached our sensory organs but instead a biased interpretation of these signals. A powerful showcase of such integration processes has been the demonstration that our perceptual decisions about current inputs are influenced by perceptual signals that immediately precede them (J. Fischer & Whitney, 2014; Ozkirli et al., 2025). For example, a previously seen line orientation or motion direction will alter the perceptual reports of consequent presentations of these features. There is much debate about what generates such serial dependence (Fritsche et al., 2017, 2020; Manassi et al., 2018; Pascucci et al., 2019), including proposals of a predictive origin (Cicchini et al., 2018). Theoretically, the prediction argument rests on humans maintaining a low estimate of environmental volatility, reflecting the spatiotemporal autocorrelations in our environment (Dong & Atick, 1995; Weilnhammer et al., 2023). Under this meta-belief of environmental stability, past inputs consequently constitute predictions about present ones and differences between the two represent the extent of prediction error, or surprise (note that we here refer to surprise as the statistical degree to which consecutive sensory states differ from one another (Schultz et al., 1997), rather than its conscious experience (Reisenzein, 2000)).

Interestingly, both the serial dependence and predictive processing literatures have sought to explain fascinating apparent conflicts in findings. Namely, that while the evidence for influences of the recent past, or predictions, on perception is compelling, the nature of that influence is far from clear. Different studies show clear evidence of attraction towards or repulsion away from recent sensory input, and other more classic predictions. What renders these conflicts especially interesting – at least in the context of prediction – is that there are good adaptive arguments for both attraction and repulsion, yet optimising for one function will come at the direct cost for another (Press et al., 2020). Attraction effects may reflect processes that amplify predicted information, to optimise accurate and efficient perception. Repulsion effects meanwhile may reflect amplification of the unpredicted information, promoting learning. If, indeed, serial dependence reflects predictive processes, then the same arguments apply also in this context (J. Fischer & Whitney, 2014; Manassi & Whitney, 2024). Optimising for speed and accuracy can be seen to therefore come at the direct cost to informativeness, and vice versa. How can this paradox be resolved?

The opposing process theory (OPT; Press et al., 2020) proposes that perception is initially attracted towards our predictions, and remains as such so long as the incoming sensory evidence does not violate them beyond what could be attributable to noise. If a sensory event does however pass this error threshold, reactive processes may lead to a gain increase in the sensory channels currently encoding the prediction-violating evidence. Such a process thus repels perception away from the invalid prediction. Importantly, this discrepancy signal is proposed to be precision-weighted so that the respective strength (or precision) of predictions and sensory evidence dictate to what extent a particular level of modal discrepancy generates a particular level of surprise (e.g., KL Divergence) and therefore elicits purported reactive processes resulting in repulsion.

Under this theory, the extent of precision-weighted discrepancy between past and current inputs should be central in determining whether perceptual decisions are attracted towards or repelled away from recent history. We here use a large amount of recently-collated trial-by-trial serial dependence data to test this idea. To pre-empt our findings, we find unanimous support for the theory across a number of analyses. When inputs are relatively similar to past ones our perceptual reports are attracted towards the past, whereas when there is a larger discrepancy they are repelled away from them. These findings thus explain a host of conflicting data from across the decades, and we consider the ways in which underlying mechanisms may provide the best optimisation for speed, accuracy and informativeness of perceptual decisions.

## Methods

### Data

We reanalyzed the trial-by-trial data compiled and standardized by Ozkirli, Chetverikov and Pascucci (2026), which collates 22 studies (Abreo et al., 2023; Blondé et al., 2023; Ceylan et al., 2021; Ceylan & Pascucci, 2023; Chetverikov & Jehee, 2023; Cicchini et al., 2018; C. Fischer et al., 2020; J. Fischer & Whitney, 2014; Fritsche et al., 2020; Fritsche & De Lange, 2019; Gallagher & Benton, 2022; Geurts et al., 2022; Houborg, Kristjánsson, et al., 2023; Houborg, Pascucci, et al., 2023; Kondo et al., 2022; Lau & Maus, 2019; Moon et al., 2023; Moon & Kwon, 2022; Ozkirli & Pascucci, 2023; Pascucci et al., 2025; Sadil et al., 2024; Samaha et al., 2019), split by within-study experiments and conditions to yield 49 samples. All studies included in the dataset employed continuous reproduction tasks. On each trial, these tasks present participants a visual display – a Gabor stimulus or a cloud of moving dots – and ask them to reproduce the orientation or motion direction (called ‘feature’ from here on) using an adjustable response bar on the screen. By considering participants’ response errors on this reproduction as a function of the previous relative to the current trial’s stimulus feature, serial dependence effects are investigated (see below). Preprocessing, including the removal of outliers, stimulus-specific biases and trials with absolute response errors or relative orientations larger than 90°, was retained from the original procedures.

### Analysis

For each trial, the response error *y* represents the signed circular difference between the presented and reported stimulus feature (positive: too far clockwise / negative: too far counterclockwise). To probe for systematic serial dependencies in these response errors, they are commonly considered as a function of *x* – the difference between the previous and the current trial’s feature. The commonly employed derivative of gaussian (DoG) model formalizes this relationship in the form:

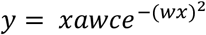

where, alongside *x* and *y, a* is the amplitude of the DoG curve peaks, *w* the curve’s inverse width and *c* the modelling constant 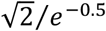 that assures that *a* numerically equals the absolute height of the curve peaks.

To model opposing processes in serial dependence data, we created a modification of this DoG model that allows the systematic direction of response errors to change along the difference between the feature on the previous and current trial – *x*. In line with our theory (Press et al., 2020), our modification allows for concurrent attraction and repulsion effects depending on surprise by modelling response errors as a weighted difference between two DoG functions (DoDoG). It takes the form:

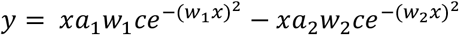

where *a*_2_ = *ratio* ∗ *a*_1_ and *w*_2_ = *w*_1_/3. The divisor of 3 was chosen to ensure a meaningfully wider, opposite signed second peak compared to the narrower first peak retained from the basic DoG model. The relative strength of the second peak was controlled through the free *ratio* parameter. Due to the conceptual implausibility – within our theory – of a narrow repulsion combined with a wide attraction effect, as well as to aid model convergence, the ratio parameter was set to 0 when *a*_*1*_ was smaller than 0.

During model-fitting, parameter bounds were equal for the two free parameters shared by the two models, *a*/*a*_*1*_ (-50;50) and *w*/*w*_*1*_ (0;0.2). The *ratio* parameter of the DoDoG model was bound between 0 and 1, ranging from no wide peak to equal amplitude narrow and wide peaks. Model-fitting was carried out by minimizing the sum of squared residuals using the R implementation of differential evolution, DEoptim (Mullen et al., 2011).

We conducted model-fitting at three levels: dataset-wide, sample-wise and participant-wise. We compared the two models using the Akaike information criterion (AIC) to account for the difference in model complexity:

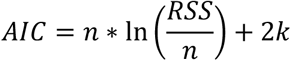

where *n* is the number of trials, *RSS* is the residual sum of squared residuals of the fitted model and *k* is the model’s number of parameters (Burnham & Anderson, 2004). Model improvement, Δ*AIC*, of our DoDoG model compared to the basic DoG model was calculated as the AIC difference between the two models: *AIC*_*DoG*_ − *AIC*_*DoDoG*_ .

### Dataset-wide analyses

We assessed the statistical significance of model parameters at the dataset-wide level using permutation-testing. Following the approach by (Fritsche & De Lange, 2019) each permutation involved randomly inverting the sign of response errors before refitting the models and retrieving model parameters. For the DoDoG model, we extracted both the a_1_ and the a_2_ parameter on each permutation. Across the 1000 permutations, this yielded permutation distributions against which we compared the empirical model parameter estimates. We applied two-sided permutation tests for all analyses.

Following recent suggestions of attractive and repulsive serial dependence effects mapping onto previous response and stimulus effects respectively, we sought to disambiguate the influences of the previous trial’s stimulus and response. Due to their considerable correlation given adequate task performance, their biasing influences on response errors are usually highly collinear. We therefore opted for a sequential regression approach where we first model the influence of one variable (e.g. previous response) and then consider the influence of the other variable (e.g. previous stimulus) on the outcome variable (response error) residuals. Due to the collinearity between the variables’ effects, this approach can be considered especially conservative since the removal of response error variance due to one variable also discards variance explained by the other, thus likely underestimating the latter’s true effect. However, the advantage is that any residual variance in the response errors can be uniquely attributed to one variable over and above the other. We conducted this approach with response first, or stimulus first, to isolate both the effect of previous stimuli and responses separately.

It is important to note that this approach is viable in the present case of an especially large dataset since the systematic underestimation of effect sizes requires considerable power to overcome.

### Participant-wise analyses

For each participant, we calculated the mean absolute error (MAE) as a proxy for sensory noise and used its reciprocal as a measure of sensory precision. We tested the participant-wise relationship between sensory precision (reciprocal MAE) and model improvement (Δ*AIC*) using afex’ R implementation for linear mixed effects regression (LMER) models (Singmann et al., 2025). Both variables were z-scored prior to LMER fitting. For all LMER analyses, we started with the maximal random effects structure and iteratively removed random effects that showed no variance or were highly correlated with other random effects (Singmann & Kellen, 2019). In this case, the resulting formula took the form:

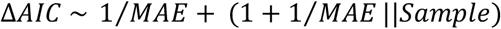

where (1 + 1/*MAE* ||*Sample*) denotes uncorrelated random slopes and intercepts for the sample to which each participant belongs.

To further characterize the link between sensory precision and effects, we extracted the absolute value of the non-zero x-intercepts of each participant’s DoDoG model fit. In other words, we recorded the surprise level at which attraction turned into repulsion. Participants showing ‘pure’ attraction or repulsion effects without an opposing, second peak in their model fit were assigned the extreme values of 0 and 90 degrees respectively. We related this surprise level to participants’ sensory precision (reciprocal MAE, both z-scored) using the LMER model:

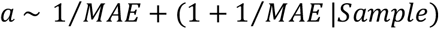

where *a* represents the absolute value of the non-zero x-intercepts of the DoDoG model fit and (1 + 1/*MAE*|*Subset*) random slopes and intercepts for the sample to which each participant belongs.

### Trial-by-trial analysis

Investigating the relationship between bias direction (attraction or repulsion) and the precision-weighted discrepancy between previous and current inputs at the trial-by-trial level, we operationalized the latter as Kullback-Leibler divergence (KLD) – given this is the formalization proposed by Press and colleagues (2020) in their opposing process theory. In simple terms, KLD quantifies the degree to which one distribution, *P*, approximates another, *Q* (Shlens, 2014). Since orientation and motion direction both occupy circular feature spaces, von Mises distributions were used to compute KLD:

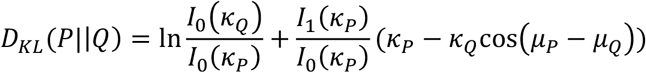

where *P* denotes the sensory evidence, *Q* the previous input and *D*_*KL*_(*P*||*Q*) the KLD between the two. *μ* and *κ* represent the respective location and concentration parameters of *P* and *Q*’s von Mises probability density function. *I*_*0*_(⋅) and *I*_*1*_(⋅) are the modified Bessel function of the first kind of order 0 and 1 respectively. For *Q* – the previous input – *μ* was the presented orientation / direction and *κ* the reciprocal absolute error on the previous trial, both in radians. For *P* – the sensory evidence – *μ* was the presented orientation / direction and *κ* the reciprocal absolute error on the current trial, both in radians. For orientation responses, the angles were multiplied by 2 before conversion to radians to map them onto the 0-360 degree space. Absolute error was again used as a proxy for sensory noise. Note that one study’s data could not be used since serial dependence was ‘induced’ by stimuli that participants did not respond to and therefore did not produce a response error (Fritsche & De Lange, 2019). Trials on which either or both of the previous and current trial’s response errors were 0 were assigned response errors of 0.001 to avoid undefined *κ* parameters (see above). Within each participant, infinite KLD values, due to very small response errors, were replaced by the highest finite KLD value. KLD was finally log-transformed to account for its positive skew.

We computed a trial-wise measure of directional response bias by inverting the sign of errors belonging to trials with negative differences between the previous and current trial feature. This is equivalent to previously used ‘folding’ procedures and allows errors to be interpreted as attracted towards (positive) or repelled away from (negative) the previous trial’s stimulus (Ozkirli et al., 2025). We related this measure of directional response bias to log-transformed KLD (both z-scored) using the following LMER model in afex:

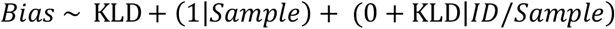

where (1|*Sample*) represents random intercepts for each sample and (0 + *KLD*|*ID*/*Sample*) represents random slopes for each participant nested within the sample to which they belong. We employed a similar model to relate KLD to the absolute size of the bias:

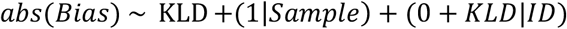

where (1|*Sample*) represents random intercepts for each sample and (0 + *KLD*|*ID*) random slopes for each participant.

## Results

We first assessed whether concurrent opposing effects generally better account for serial dependence data than exclusively attractive or repulsive effects by comparing the two models on data pooled over all participants and studies. In line with previous research, the basic DoG model showed an attraction effect (*a* = 1.569) that peaked when previous and current trial stimuli were 18.532° apart (*w* = 0.038). The DoDoG similarly accounted for the narrow attraction effect (*a*_1_ = 1.679, *p* = 0.005; *w*_1_ = 0.034), however concurrently also captured a wide repulsion (*ratio* = 0.149; *a*_2_ = 0.250, *p* = 0.013; *w*_2_ = 0.011). Despite the added complexity in the form of an extra free parameter, the DoDoG model nevertheless strongly outperformed the basic DoG model, Δ*AIC* = 59.067 (Figure 2A). The DoDoG curve thus suggests that perceptual decisions are attracted towards recent history when the discrepancy is low and repelled away when it is large.

**Figure 1.**
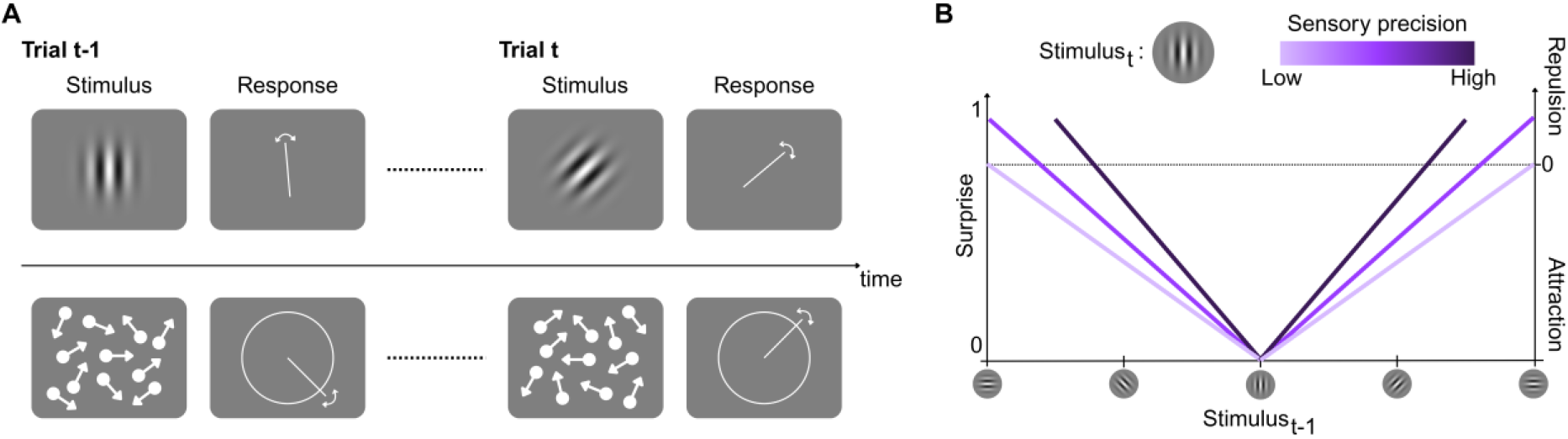
**A** Schematic illustration of the serial dependence paradigms included in the analysed dataset (top: oriented Gabor, bottom: dot motion clouds). On each trial, participants see and then report a visual stimulus feature using a continuous adjustment response. Note that studies frequently employ backwards masks and delays between stimuli and responses, which are omitted in this schematic. Serial dependence biases constitute systematic response errors on trial t as a function of the stimulus (and/or response) on trial t-1. **B** Conceptual illustration of predicted serial dependence biases under the opposing process theory, here illustrated for oriented Gabors. Given a belief of environmental stability, a stimulus on trial t evokes different levels of surprise depending on its modal discrepancy with respect to the stimulus on trial t-1. Serial dependence biases are predicted to vary systematically with surprise. When surprise is low, serial dependence is attractive. When surprise is especially high (passing threshold signalled with a dashed line), serial dependence turns repulsive. Sensory precision scales the mapping of modal discrepancy (x-axis) onto surprise (left y-axis), and therefore also accelerates the shift from attraction to repulsion (right y-axis).

**Figure 2.**
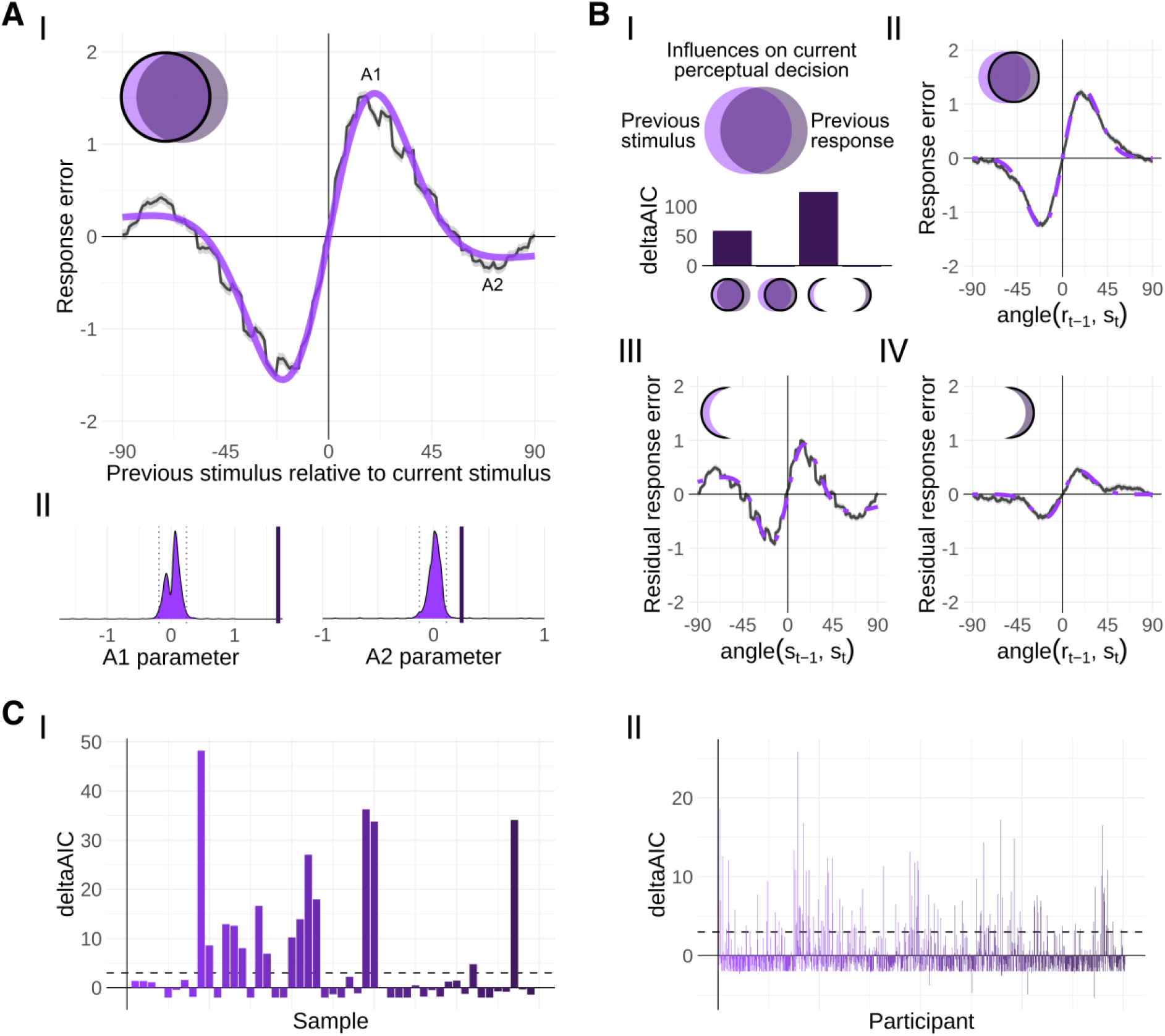
**A** Dataset-wise effects of previous stimuli on response errors. (I) Solid black line and shading represent grand-averaged sliding mean and standard error of mean. Purple line represents DoDoG model fit. A1 and A2 are the model parameters reflecting the size of narrow attraction and wide repulsion effects respectively. (II) Permutation distributions and empirical parameter estimates for DoDoG fit shown above. Solid vertical lines represent the empirical model parameters and dotted vertical lines the significance threshold for two-sided permutation tests. **B** (I) Top: Schematic illustration of the collinear influences of previous stimuli and responses on current perceptual decision-making. Bottom: Model improvement (deltaAIC) of DoDoG over DoG for variance-partitioned data displayed in A and remainder of B. (II) Effect of previous responses on response errors. (III) Effect of previous stimuli on response-cleaned residual response errors. (IV) Effect of previous responses on stimulus-cleaned residual response errors. **C** (I) Sample-wise model improvement (deltaAIC) of DoDoG over DoG. (II) Same as (I) but for each participant. Participant-wise colours match colours in (I) and denote sample membership.

We next conducted analyses to shed light on the relative contributions of decisional or sensory history. This approach follows recent proposals that attraction and repulsion effects map onto perceptual and decisional origins respectively (Fritsche et al., 2020; Pascucci et al., 2019). Given a sufficient level of task performance, previously presented stimuli and responses should correlate strongly, and this is indeed the case in this dataset, *r* = 0.887. This collinearity renders the respective influences more difficult to discern (Figure 2B-I), but not impossible with a dataset of this size. Using a sequential regression approach (see Methods), we found that while the effect due to previous responses was best described by a purely attractive model: *a*_1_ = 0.432, *p* = 0.010; *w*_1_ = 0.040; *ratio* = 0, Δ*AIC* = −2 (Figure 2B-IV, non-residual: 1B-II), the effect due to previous stimuli reflected the bidirectional pattern of attraction turning into repulsion with increasing discrepancy: *a*_1_ = 1.096, *p* = 0.008; *w*_1_ = 0.039; *ratio* = 0.308; *a*_2_ = 0.338, *p* = 0.013; *w*_2_ = 0.013, Δ*AIC* = 124.346 (Figure B-III).

At the sample and participant-level, we observed considerable variability in the extent to which the DoDoG outperformed the DoG model (Figure 2C). We therefore asked whether this variability reflected parameters hypothesized to determine it under our model. Specifically, we considered whether higher precision of sensory evidence determines that it is more likely the DoDoG will outperform the DoG model, because there is a higher chance that signals will have been sufficiently discrepant to elicit the purported reactive processes (Press et al., 2020). To approximate sensory precision in each participant, we here used their behavioural performance, inversely represented in their mean absolute error (MAE). Logically, participants should perform better if sensory evidence is precise and worse if it is imprecise. Recent neuroimaging research additionally supports the assumption that behavioural errors reflect increased neural noise (Geurts et al., 2022). Note that various factors could influence the precision of sensory evidence, including study-specific stimulus parameters and participant-specific capacities, but they are importantly all reflected in performance. After fitting both models to each participant individually, we first predicted model improvement (Δ*AIC*) from sensory precision (reciprocal MAE). Since the two models are nested (ratio = 0 renders DoDoG equal to DoG), any model improvement could be attributed to the presence of high-discrepancy repulsion effects for which the basic DoG model cannot account. Interestingly, sensory precision and model improvement of DoDoG over DoG indeed showed a positive relationship, *β* = 0.134, *t*(28.558) = 3.272, *p* = 0.003, *d* = 1.22 (Figure 3A), in line with the theory’s prediction.

**Figure 3.**
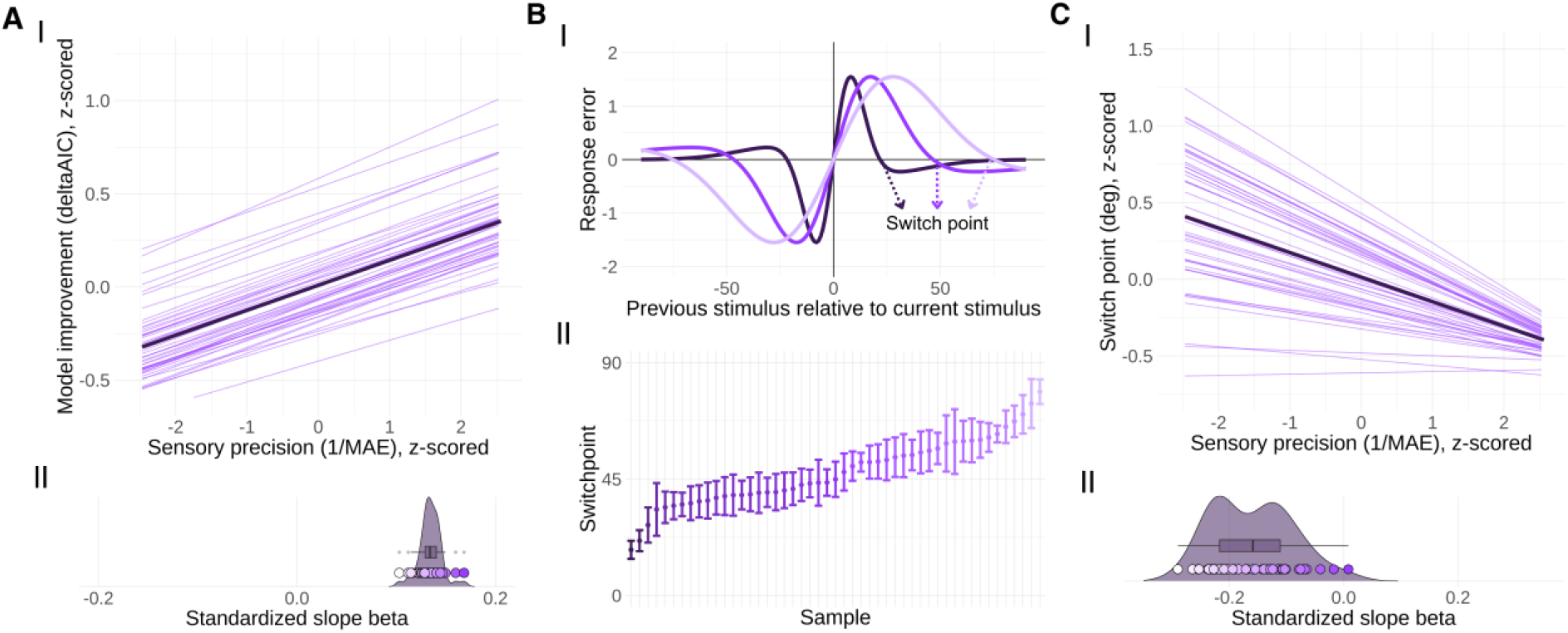
**A** Participant-wise relationship between sensory precision and model improvement (deltaAIC) of DoDoG over DoG. (I) Each thin line represents a sample displayed using its random slope and intercept from the linear mixed effects model fit to participant-level data. The thick, darker line represents the model’s fixed effect. (II) Linear mixed effects model random slopes for each sample. Each circle represents a sample and its colour the strength of the effect. The boxplot encompasses the interquartile range (IQR) around the median and its whiskers extend to 1.5*IQR below and above the first and third quartile respectively. **B** (I) Schematic illustration of the switch point, with the non-zero x-intercept of the DoDoG curve reflecting the point at which attraction turns into repulsion. From light to dark hues the switch point moves closer to 0, modelled as reflecting surprise at lower discrepancy levels due to increased sensory precision. (II) Switch point for each sample. Points represent the sample mean and the error bars reflect its standard error. **C** Participant-wise relationship between sensory precision and switch point width. Same plotting conventions apply for (I) and (II) as in **A**.

Small modal differences between predictions and sensory evidence would elicit the same levels of surprise under high sensory precision, as larger modal differences under low precision – deemed functionally adaptive because small differences are less likely attributable to noise in the former case (Press et al., 2020). If the switch from attraction to repulsion effects is a function of surprise, for higher sensory precision we would thus expect the switch to occur at smaller modal differences between predictions and sensory evidence (Figure 3B). At the extremes, ‘pure’ attraction and repulsion in the absence of a cooccurring effect in the opposite direction would thus result from particularly low and high sensory precision, respectively. We indeed found a negative relationship between sensory precision (reciprocal MAE) and the switch point at which response biases flipped from attraction to repulsion, *β* = −0.160, *t*(14.388) = −3.841, *p* = 0.002, *d* = 2.03 (Figure 3C). This combination of findings importantly suggests that sensory precision amplifies prediction-evidence-mismatches and therefore reduces the modal error threshold required to generate repulsion effects.

Having established the link between surprise and direction of effect in individual participants, we next sought a more fine-grained approach in the form of a trial-by-trial analysis of discrepancy. We computed Kullback-Leibler divergence (KLD) on each trial by treating the previous trial’s stimulus as the prior distribution for the current trial and comparing it against the input distribution of the current trial (Figure 4A, see Methods). In short, precision of prior and input were again approximated using the response error while their modes represented the previously and currently presented feature respectively. KLD therefore represented the trial-by-trial precision-weighted discrepancy between previous and current stimuli. We related KLD to a trial-by-trial, directional bias measure: the number of degrees that the response error was attracted towards (positive) or repelled away (negative) from the previous trial’s feature. Note that this equates to inverting the sign on response errors in the two left-hand quadrants in Figure 2A, in line with common folding procedures (Ozkirli et al., 2025). We first found that absolute biases were larger when KLD was smaller (*β* = −0.252, *t*(927.045) = −60.56, *p* < 0.001, *d* = 3.98). This is unsurprising given that KLD is partly based on distributional precision which we tethered to response errors. More interestingly than their magnitude, the directionality of the response biases also changed with surprise. Specifically, we found that higher KLD was associated with more repulsion (*β* = −0.004, *t*(511.910) = −2.438, *p* = 0.015, *d* = 0.22, Figure 4B&C). We were thus able to show that serial dependence biases turn from attractive to repulsive depending on surprise across the sample, participant, and trial level of this dataset.

**Figure 4.**
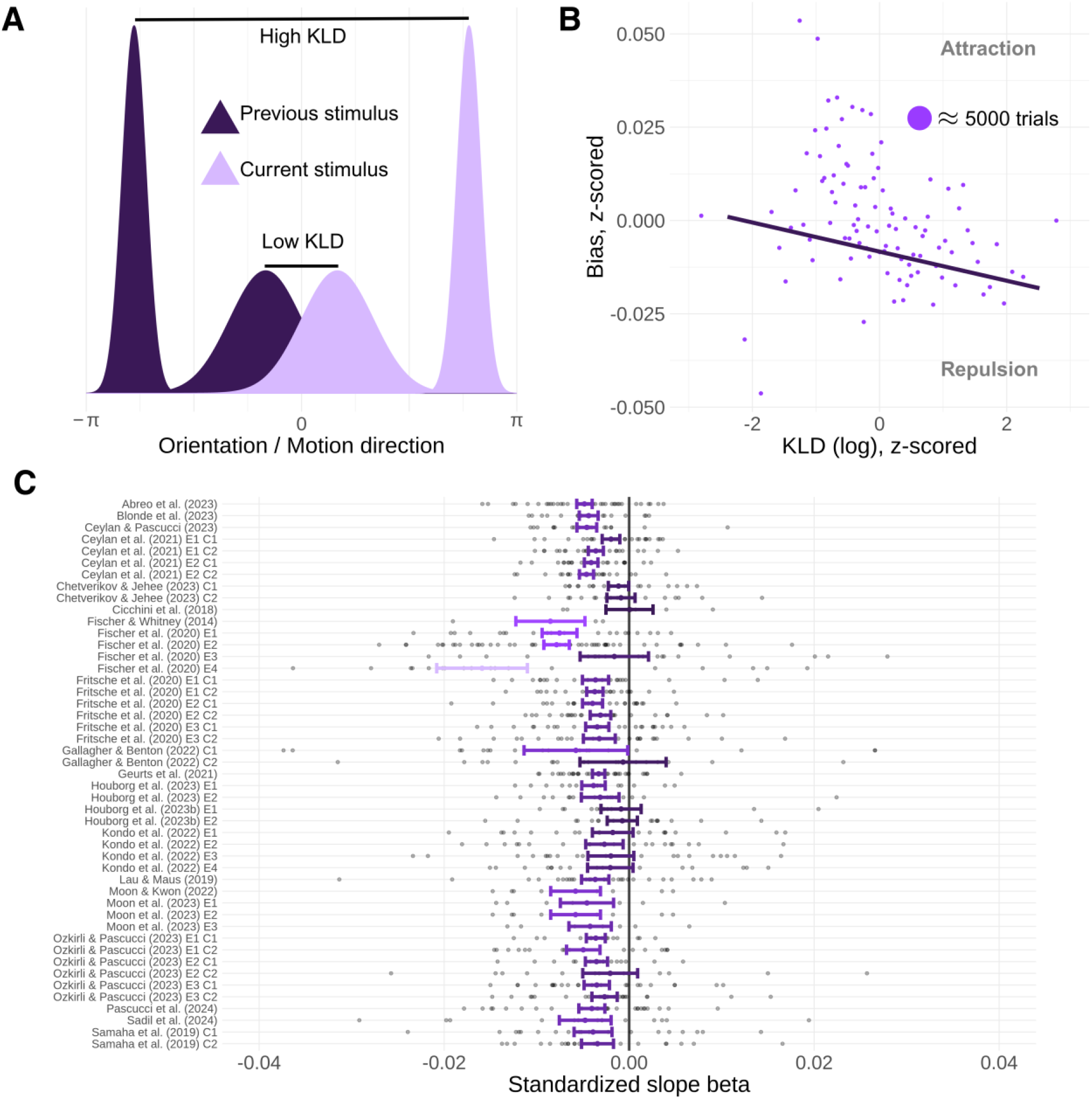
**A** Schematic illustration of Kullback-Leibler divergence (KLD) as implemented on serial dependence trial-by-trial data. Note that the logic is here displayed in Cartesian dimensions using Gaussian distributions but is implemented using circular von Mises distributions during modelling. The feature (orientation / motion direction) space is expressed in radians to incorporate the different possible stimulus values in orientation (-90 to 90) and motion direction (-180 to 180). **B** Trial-wise relationship between KLD and Bias. The thick, darker line represents the fixed effect from the linear mixed effects model. For display purposes only, we collapsed over participants and sorted trials by KLD before evenly dividing them into 100 bins of approximately 5000 trials each. **C** Random slopes for each participant from the linear mixed effects model, grouped by sample. Within each sample, each grey circle represents a participant, the coloured point the sample-wise random slope mean and the error bars its standard error. Sample names are retained from Ozkirli, Chetverikov & Pascucci (2026). The consistently negative slope beta suggests that biases flip from attractive to repulsive with increasing KLD.

## Discussion

Recent perceptual inputs bias perceptual decision-making but when, why, and how has remained a topic of debate. We here leveraged a large-scale trial-by-trial dataset to show that serial dependence biases can be considered as a function of the precision-weighted discrepancy between previous and current sensory inputs. Biases towards past inputs (‘attraction’) gave way to biases away from them (‘repulsion’) as this discrepancy increased. More precise sensory evidence accelerated the shift from attraction to repulsion, in line with a precision-weighted inference process. Modelling this precision-weighted discrepancy as KL divergence, we finally showed that it explains both the strength and direction of serial dependence effects on a trial-by-trial basis.

These findings provide broad support for the opposing process theory, which proposes that a surprise-moderated shift from attraction to repulsion would be adaptive (Press et al., 2020). Given that we live in a rather stable environment (my laptop is more likely to remain a laptop than turn into a typewriter from one moment to the next), attraction towards previous sensory states allows perception to remain efficient and veridical in the face of sensory noise. However, when we collect strong evidence that current sensory states no longer reflect past ones, repulsion allows us to update our world model and register this change in our environment. Interestingly, the opposite argument appears to have been made recently within the serial dependence literature – that these history-dependent biases have an overall detrimental effect on perceptual decision-making (Ozkirli et al., 2025). However, as Ozkirli and colleagues note, serial dependence paradigms, by design, contain temporally uncorrelated stimuli, while our natural environment is filled with temporal autocorrelations that do make the current sensory environment a capable predictor of future sensory states. Attractive ‘continuity fields’ may thus exploit this structure by cutting through uninformative noise, while high surprise repulsion assures that true changes in the environment are not concealed (Manassi & Whitney, 2024). We therefore argue that the mechanisms yielding attraction optimise accuracy on average in our natural environments, even if lab settings intentionally disrupt natural structure and thus do not showcase these accuracy benefits.

High-discrepancy repulsion accompanying low-discrepancy attraction, though previously noted on several occasions (Fritsche & De Lange, 2019; Gallagher & Benton, 2022; Houborg, Pascucci, et al., 2023; Rafiei, Chetverikov, et al., 2021; Rafiei, Hansmann-Roth, et al., 2021; Samaha et al., 2019; Van Bergen & Jehee, 2019), is not uniformly apparent in serial dependence studies and thus has led some to doubt its consistency (Manassi et al., 2023). Our findings suggest that this apparent inconsistency is precisely expected given study and participant-wise differences in the surprise-related variables governing the flip from attraction to repulsion. Describing this flip in effect direction as a function of surprise (KLD) establishes clear roles for contextual variables like sensory precision evidence that had previously remained unclear (Ceylan et al., 2021; Gallagher & Benton, 2022; Van Bergen & Jehee, 2019). For example, the reported reduction in (attractive) serial dependence due to reduced sensory uncertainty can be understood as a shift from attraction to repulsion effects that is not captured by one-directional models of serial dependence (Ceylan et al., 2021; Cicchini et al., 2018; Gallagher & Benton, 2022). More generally, the success of a predictive processing-inspired theory in explaining serial dependence arguably gives some credence to past proposals of predictive processes underlying these effects (Burr & Cicchini, 2014; Kalm & Norris, 2018).

Notably, even if prediction mechanisms contribute to serial dependence, other mechanisms will also influence processing and their ability to explain these effects should be considered. Most plausibly, previous explanations of repulsive serial dependence have frequently pointed to sensory adaptation as playing an important role (Fritsche et al., 2020; Moon & Kwon, 2022; Pascucci et al., 2019). Specifically, stimulation of a particular neural subpopulation temporarily reduces its responsiveness to subsequent inputs and thereby biases consequent percepts away from recent ones (Thompson & Burr, 2009). Serial dependence likely reflects a combined influence of adaptation and other predictive processes (Fritsche et al., 2020). Nevertheless, it is unclear how adaptation mechanisms could generate the here-observed flip from attraction to repulsion dependent upon surprise. In fact, adaptation-generated repulsion would be more likely at smaller discrepancies, given that nearby stimuli engage maximally overlapping neural populations (Clifford et al., 2000). Furthermore, the narrow tuning of sensory adaptation has even been found to turn attractive at high discrepancy values, thus reflecting the opposite pattern to that reported here (Clifford, 2014; Clifford et al., 2000). Indeed, a previous model of concurrent attractive and repulsive serial dependence effects, modelling repulsion using efficient coding, only scarcely captured wide repulsion effects and the same authors conceded elsewhere that sensory adaptation was unlikely to account for these effects (Fritsche et al., 2020; Fritsche & De Lange, 2019). Nevertheless, consideration of how these different types of mechanisms might dissociate, or indeed interact, remains an important task for future research.

Relatedly, we here found a purely attractive effect of previous responses on perceptual decisions, alongside the more nuanced influence of previous sensory events. The response effects are in line with previous proposals of decisional inertia or persevering working memory templates leading to attractive effects (Fritsche et al., 2020; Moon & Kwon, 2022; Pascucci et al., 2019). However, the effect of previous stimuli perhaps contradicts those proposals’ reports of purely repulsive influences (although see Cicchini et al., 2017, 2018; Manassi et al., 2018). In addressing this discrepancy, it is important to note that while dissociating the stimulus and response origins of previous trial influences, our analysis cannot speak towards the level at which these influences affect perceptual decision-making on the current trial. These may, at least in theory, still have occurred at a purely decisional level. Although our analyses were inspired by a theory that does frame attractive predictive influences as perceptual (Press et al., 2020), serial dependence likely also involves repulsive perceptual adaptation effects at low discrepancy levels (although see: Janetsky et al., 2026) and so the surprised-mediated flip from attraction to repulsion may instead occur at a post-perceptual stage of processing. Whether serial dependence effects occur at the perceptual or decisional level thus remains an open question, as evidenced by the considerable heterogeneity in the literature to this day.

Finally, it is worth highlighting that our findings imply a plausible explanation for null findings of predictive effects – here in the form of serial dependence. Specifically, we found a level of precision-weighted discrepancy between prediction and input where influences change from attractive to repulsive. Experimental manipulations sampling from such intermediate levels of surprise – whether within or across participants – should therefore be expected to produce null findings, particularly in binary response paradigms that do not map out a gradient of surprise. Our findings thus highlight the value of the precedent set by serial dependence research of employing continuous reproduction tasks when studying prediction effects on perceptual decision-making, as some have already done (Sánchez-Fuenzalida et al., 2023). In turn, we also highlight the value in formalizing a theory that predicts *when* effects will be attractive, repulsive, or null, rather than simply dismissing nulls as potentially hitting this switch point.

Perception has been paradoxically shown to be attracted towards and repelled away from recent sensory inputs. Drawing on recent theories of predictive influences on perception, we show that the direction of this biasing influence can be predicted based on the precision-weighted discrepancy between past and current sensory input. Functionally, this may allow perception to remain efficient while allowing for the detection of change between the past and present. We showcase the reliability of this relationship by finding support for it across a large-scale dataset.

## Acknowledgements

This work was funded by a Leverhulme Trust project grant (RPG-2022-358), and European Research Council (ERC) consolidator grant (101001592) under the European Union’s Horizon 2020 research and innovation programme, both awarded to CP. We are grateful to other members of the Action and Perception Lab (especially Kirsten Rittershofer, Nicholas Simpson, Quirin Gehmacher, Jessye Clarke, Gamze Bilgen and Richard O’Farrell), Peter Kok, and Floris de Lange, for useful discussions.

